# Three-dimensional analysis and *in vivo* imaging for sperm release and transport in the murine seminiferous tubule

**DOI:** 10.1101/2022.01.31.478089

**Authors:** Yuta Kanazawa, Takuya Omotehara, Hiroki Nakata, Tsuyoshi Hirashima, Masahiro Itoh

**Affiliations:** Department of Anatomy, Tokyo Medical University, Tokyo, Japan; Department of Histology and Cell Biology, Graduate School of Medical Sciences, Kanazawa University, Kanazawa, Japan; The Hakubi Center, Kyoto University, Kyoto, Japan; Department of Pathology and Biology of Diseases, Graduate School of Biostudies, Kyoto University, Kyoto, Japan; Japan Science and Technology Agency, PRESTO, Kawaguchi, Japan

**Keywords:** *In vivo* imaging, Seminiferous tubule, Spermatozoa, Three-dimensional analysis

## Abstract

**Introduction:** Spermatozoa released from Sertoli cells must be transported to the epididymis. However, the contribution of the peristaltic motion in the seminiferous tubule to sperm release and transport remains unclear. We, therefore, investigated luminal flow and movements in the seminiferous tubules by three-dimensional analysis and *in vivo* imaging.

**Materials and Methods:** Serial testicular sections were cut in 5-μm-thick and 50-μm-interval and stained by PAS-hematoxylin. After the three-dimensional reconstruction of the seminiferous tubules, the localization of the flowing spermatozoa and stages observed in the sections were recorded in each reconstructed tubule. The luminal movements in the seminiferous tubule were observed by *in vivo* imaging using a fluorescent-reporter mouse and two-photon excitation microscopy system.

**Results:** Flowing spermatozoa were mainly scattered in the lumina at stage VII/VIII, and clustered spermatozoa were also found in some regions. The clustered spermatozoa were observed at zero to two regions in each seminiferous tubule. Flowing spermatozoa were also found in the opposite direction to the rete testis. The flagellum direction of the spermatozoa attached to the seminiferous epithelium was reversed within a few seconds to a few tens of seconds when observed by *in vivo* imaging. The epithelium at the inner curve of the seminiferous tubule moved more actively and attached fewer spermatozoa compared to that at the outer curve.

**Discussion:** This study revealed the presence of repeatedly reversed luminal flow in the seminiferous tubule. Such movements are suggested to help the sperm release from the Sertoli cells and the following aggregation of the released spermatozoa.

## Introduction

Spermatozoa differentiate in the seminiferous tubule (ST) and are released into its lumen. Because the released spermatozoa possess no fertility, they must be transported to the epididymis, where the spermatozoa acquire motility and capacitation. Before the transport to the epididymis, spermatozoa flow through the ST, tubuli recti, rete testis, and efferent duct of testis. Therefore, the ST is the start of the male genital tract. Daily sperm production in mammalian species is 4–6×10^7^ per gram of testis (Hess, 2008). Because the testis weight is approximately 0.1 g in some mouse strains (Suto, 2008), about 4–6×10^6^ spermatozoa should pass through the STs toward the rete testis per day in a mouse testis.

Spermatozoa do not possess active motility in the ST (Holstein *et al*., 2003). Additionally, a recent study revealed that Neural Epidermal Growth Factor–like–like 2 (NELL2) secreted from testicular germ cells induces the differentiation of the downstream of the male genital tract, the initial segment of the epididymis, which is necessary for sperm maturation (Kiyozumi *et al*., 2020). Therefore, some mechanisms to transport the spermatozoa and luminal fluid from the seminiferous tubule to the epididymis are needed. Previous studies have focused on the peritubular myoid cells. The transport of the spermatozoa is induced by the peristaltic motion of the ST to approach the epididymis (Maekawa *et al*., 1996). Maekawa et al. (1996) reported that “the frequency of the pressure change in the seminiferous tubule is 0.17-0.23Hz, and is lower than that of epididymal ducts where several muscle layers and rich innervation are present.” The diameter of the rat STs shows a 40% reduction when they contract (Losinno *et al*., 2016). When the function of the peritubular myoid cell (PMC) is disrupted, the luminal flow in the ST is modulated, resulting in dilation of the ST (Uchida *et al*., 2020). Most recently, unidirectional movement of the spermatozoa transported in the lumina of the ST was shown by an *in vivo* imaging method (Fleck *et al*., 2021). Motile cilia on the epithelial cells of the efferent tubules are also found to generate luminal flow using *in vivo* imaging (Yuan *et al*., 2019). Additionally, more than 90% of the luminal liquid from the testis is absorbed between the rete testis and efferent tubules (Clulow *et al*., 1994), and ligation of the efferent duct induces the dilation of the ST and rete testis (Anton, 1979). Therefore, the liquid absorbance also contributes to the luminal flow. Although the transport of the spermatozoa was recently reported as above, the luminal flow throughout a whole ST cannot be observed by *in vivo* imaging. Furthermore, we wondered how the spermatozoa were clustered in the ST after release from the Sertoli cells.

Spermatids are transformed into mature spermatozoa via 16 and 19 steps called spermiogenesis in mice and rats, respectively (Russell *et al*., 1990; Nakata *et al*., 2015a). During the steps, they need to be attached to the Sertoli cells by some adhesion structures, such as the ectoplasmic specialization and tubulolobular complex (Lie *et al*., 2010). Sertoli cells also need to form certain cytoskeleton networks inside them (O’Donnell *et al*., 2011; Yan Cheng and Mruk, 2015). These structures and molecules, including a focal adhesion kinase, are tightly regulated during spermiogenesis, and therefore, sperm release called spermiation is induced via these regulations. Disruption of the regulation leads to the failure of spermiation, resulting in the persistence of the mature spermatozoa in the seminiferous epithelium even after stage IX. In contrast, when the adherent molecule in the ectoplasmic specialization, a testis-specific adherens junction, is deleted in the mouse, the spermatids after the specific step are reduced on the seminiferous epithelium (Nakata *et al*., 2015a).

The three-dimensional (3D) reconstruction method is helpful for us to investigate the whole structure of the STs and has revealed the 3D structure of the STs, including the distribution of stages of the seminiferous epithelium (Nakata *et al*., 2017). Furthermore, a high dose of busulfan induces impairment of the spermatogenesis in the longer serial ST region, instead of the more number of the impaired region (Nakata *et al*., 2020). Here, to estimate the sperm flow from detaching out of the Sertoli cell to reaching the rete testis, we investigated the localization of the flowing spermatozoa in each ST using a 3D reconstruction model. Furthermore, the luminal movement in the ST was observed by *in vivo* imaging using a fluorescent-reporter mouse and two-photon excitation microscopy system.

## Materials and methods

### Animal

The present animal study for three-dimensional analysis was approved by Kanazawa University (approval number: AP-153636) and conducted in accordance with the Guidelines for the Care and Use of Laboratory Animals in Kanazawa University. C57BL/6 strain male mice were purchased from Nippon SLC, Inc., reared under standard 12-h light/12-h dark laboratory conditions with free access to standard food and water, and used at the age of 90 days.

Pax2-LynVenus mouse was produced by crossing the Pax2-Cre mouse (Ohyama and Groves, 2004) and R26R-Lyn-Venus mouse (Abe *et al*., 2011) at Kyoto University. All the animal experiments for *in vivo* imaging were approved by the local ethical committee for animal experimentation (MedKyo 19090 and 20081) and were performed in compliance with the guide for the care and use of laboratory animals at Kyoto University. For histological analysis using Pax2-LynVenus mice, they were maintained at a specific pathogen-free facility in Tokyo Medical University. They were maintained at 22–24 deg C and 50%–60% relative humidity with a 12-hours light-dark cycle. This study was approved by the Tokyo Medical University Committee (Permission #R2-0042).

### Histological analysis

C57BL/6 strain male mice were sacrificed by cervical dislocation. The testis and epididymis were dissected out *en bloc*, fixed by immersion in Bouin’s solution overnight, dehydrated in a graded ethanol series, and embedded in paraffin. Serial 5-μm-thick sections with intervals of 50 μm were made from three testes by cutting the specimen longitudinally in parallel to the plane involving both the testis and epididymis using a microtome, and then mounted on glass slides. The sections were treated with Periodic acid-Schiff-hematoxylin (PAS-H) to stain the basement membrane of seminiferous tubules, as previously described (Nakata and Iseki, 2019). The sections were digitized using a whole-slide scanner (Nanozoomer 2.0-HT; Hamamatsu Photonics) with a 20-fold objective lens. The resulting digital images of the sections were visualized with viewer software (NDP.view2; Hamamatsu Photonics). The number of ST’s cross sections, the number and position of spermatozoa apart from the seminiferous epithelium in the ST, and the stage of the seminiferous epithelium was recorded throughout the captured pictures.

Pax2-LynVenus male mice were anesthetized using 2% isoflurane and euthanized by cervical dislocation. The testis and epididymis were collected and flash-frozen in O.C.T. compound (Sakura Finetek Japan) with liquid nitrogen. The frozen block was cut in 5-μm-thick using Cryostar NX70 (Thermo Fisher Scientific) and fixed in 10% formalin in 0.1 M phosphate buffer pH7.4 for 10 min at room temperature. After being washed in 0.1 M phosphate buffered saline pH7.4 containing 0.5% Tween 20 (PBST), the sections were incubated with peanut lectin (PNA) conjugated to Alexa Fluor 647 (L32460, Thermo Fisher Scientific, 1:2,000) and DAPI (340-07971, DOJINDO, 1:1,000) for 1 hr at room temperature. The PNA was stained for staging the STs (Nakata *et al*., 2015a). After PBST washes, they were coverslipped in FluorSave (Merck Millipore). The observation was performed using LSM700 (Zeiss).

### Three-dimensional analysis

The 3D reconstruction was performed as previously described (Nakata *et al*., 2017). Briefly, the digital images of serial sections were extracted from the PAS-H-stained basement membrane representing the outline of seminiferous tubules and then converted into grayscale in JPEG format using Adobe Photoshop 2020 software (Adobe Systems). Using Amira 6.3.0 software (Visage Imaging), the inside of the basement membrane of the selected tubule was filled with a particular color using threshold processing and traced from section to section. This procedure was repeatedly applied to all STs, and they were then 3D-reconstructed. The core lines of all reconstructed STs were also drawn with the same software, and the number of the STs was recorded from the cranial side. The counted spermatozoa were pointed on the core line, and the distance from the nearest rete testis was recorded. The core lines were segmented according to the stage of the seminiferous epithelium divided into three groups, stage I-VI, VII/VIII, and IX-XII.

### *In vivo* imaging

For intravital imaging, male mice were anesthetized with vaporized isoflurane at 1.5% by the inhalation device (Biomachinery, TK-7). An incision in the murine scrotum and the parietal layer of tunica vaginalis was made by surgical scissors, and the testis was gently pulled out. The mouse was laid on an electric heat pad (#MATS-U52AXKA26-A13L, Tokai Hit) maintained at 37°C in the prone position, and the testis was set on a glass coverslip (#C024501, Matsunami Glass Ind.) attached to the heat pad for inverted observation. Then, two-photon excitation microscopy was performed with an inverted microscope system (FV1200MPE-IX83, Olympus), equipped with the ×30 silicone-immersion lens (NA=1.05, WD=0.8 mm, UPLSAPO30XS, Olympus). Time-lapse images were acquired at 945 nm excitation wavelength (InSight DeepSee, Spectra-Physics) with the emission filter (BA520-560, Olympus). The STs at stage VII/VIII were observed because of the presence of the sperm flagella.

## Results

### The number of spermatozoa in the sections

We counted over 10,000 cross sections of the ST (Table 1). Spermatozoa apart from the seminiferous epithelium were counted as the flowing spermatozoa in the lumen (Fig. 1a-e). As a result, the ratio of the ST cross sections containing the flowing spermatozoa was approximately only 5% (Table 1). The flowing spermatozoa were mainly scattered in the ST lumen (Fig. 1b-d). A previous study reported that the flowing spermatozoa were observed as a cluster through the ST by *in vivo* imaging in the bright field (Fleck *et al*., 2021). Therefore, we tried to find the clustered spermatozoa in the lumen and found that some regions contained them (Fig. 1e). In Total, 1–3×10^3^ flowing spermatozoa were observed (Table 1), and therefore, the number of spermatozoa in a whole testis was estimated to be around 1–3×10^4^ by multiplying 10. Additionally, when focusing on the ST structure throughout the cross sections, we found characteristic attachment of the spermatozoa on the seminiferous epithelium, especially in the curved STs. As the previous three-dimensional analysis on the STs revealed (Nakata *et al*., 2015b, 2017), the STs are well twisted. Therefore, some longitudinal cross sections of the curved STs can be observed. There were less than a few curved STs at stage VII/VIII when the most mature spermatozoa were observed in a series of whole testis cross sections. We found that fewer spermatozoa were attached to the top of the seminiferous epithelium at the inner curve than that of the outer curve (Fig. 1f,f’). Spermatids at step 8, which usually localize beneath the spermatozoa at step 16 at stage VIII, were observed even at the top of the epithelium of the inner curve.

**Figure 1.**
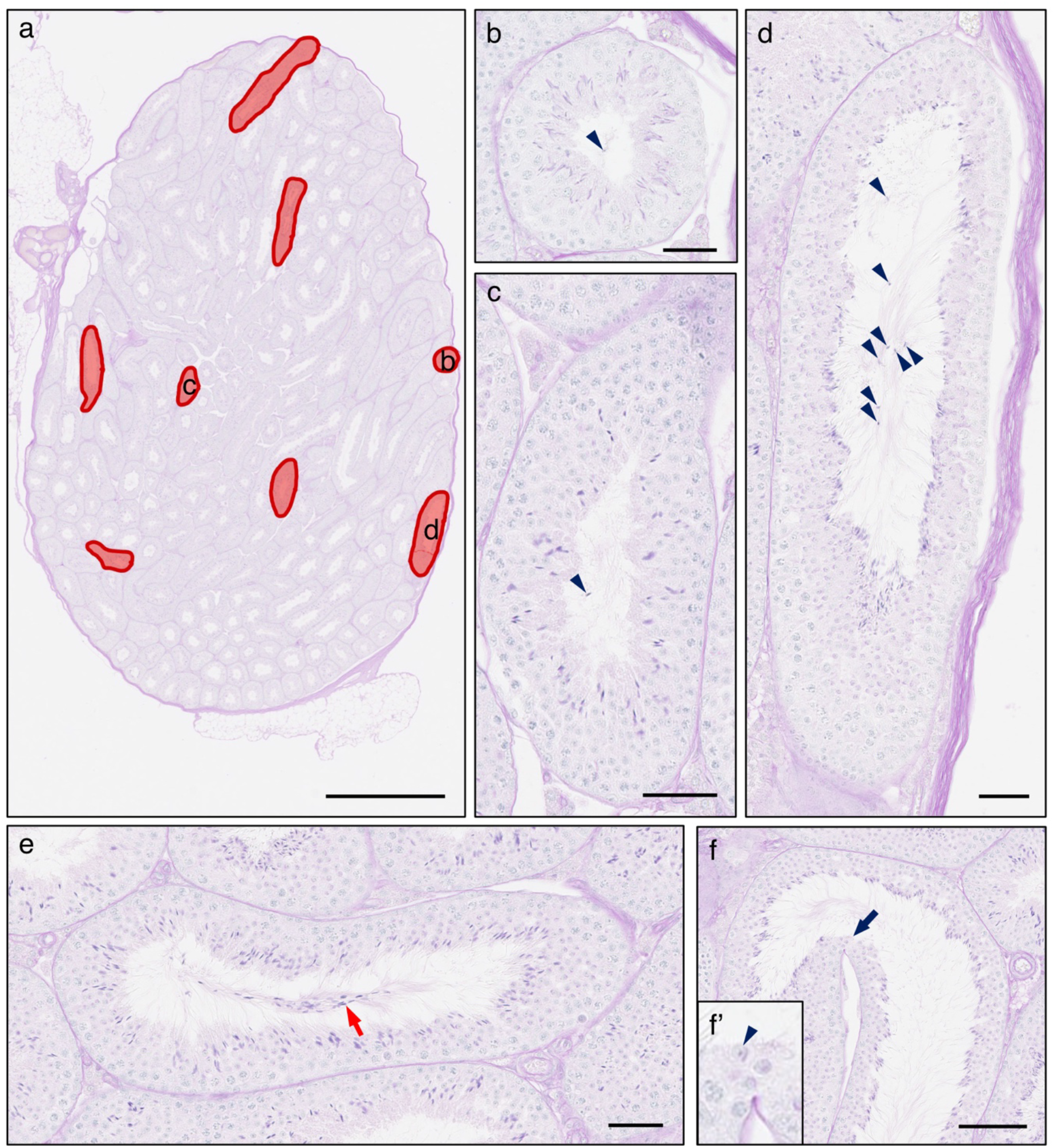
Representative pictures of flowing spermatozoa and seminiferous tubules in testicular sections. (a-d) A few seminiferous tubules containing flowing spermatozoa (red circles) are observed in a section (a), and their magnified pictures are shown in b-d. Arrowheads indicate flowing spermatozoa. (e) Clustered spermatozoa (a red arrow) in another seminiferous tubule are also observed. (f,f’) Fewer spermatozoa are attached to the seminiferous epithelium at the inner curve in the seminiferous tubule (a black arrow), but there are spermatids at step 8 (arrowhead). Bars = 1 mm (a), 50 μm (b-e), 100 μm (f).

**Table 1.** The number of counted seminiferous tubules and flowing spermatozoa in each testis

|  | Counted ST<br>(A) | ST containing<br>flowing sperm<br>(B) | Ratio of ST containing<br>flowing sperm<br>B/A *100 (%) | Sperm count |
| --- | --- | --- | --- | --- |
| #1 | 10,776 | 481 | 4.46% | 1,869 |
| (stage VII/VIII) | (1,285) | (263) | (20.5%) | (1,142) |
| #2 | 10,772 | 537 | 4.99% | 1,687 |
| (stage VII/VIII) | (1,467) | (217) | (14.8%) | (895) |
| #3 | 13,120 | 835 | 6.36% | 2,856 |
| (stage VII/VIII) | (2,563) | (547) | (21.3%) | (1,941) |

### The number of spermatozoa in an intact seminiferous tubule

We next investigated the number and localization of the flowing spermatozoa in each intact ST using the 3D reconstruction model. After all STs were reconstructed, three simple STs with no or fewer branching points were chosen from the cranial, middle, and caudal regions from three testes. The localization of the flowing spermatozoa was plotted on the reconstructed ST (Fig. 2a). There were some regions where the flowing spermatozoa were frequently observed. Such areas were not biased by the cranial-caudal or dorsal-ventral axis. Therefore, the stage of the STs was next recorded.

**Figure 2.**
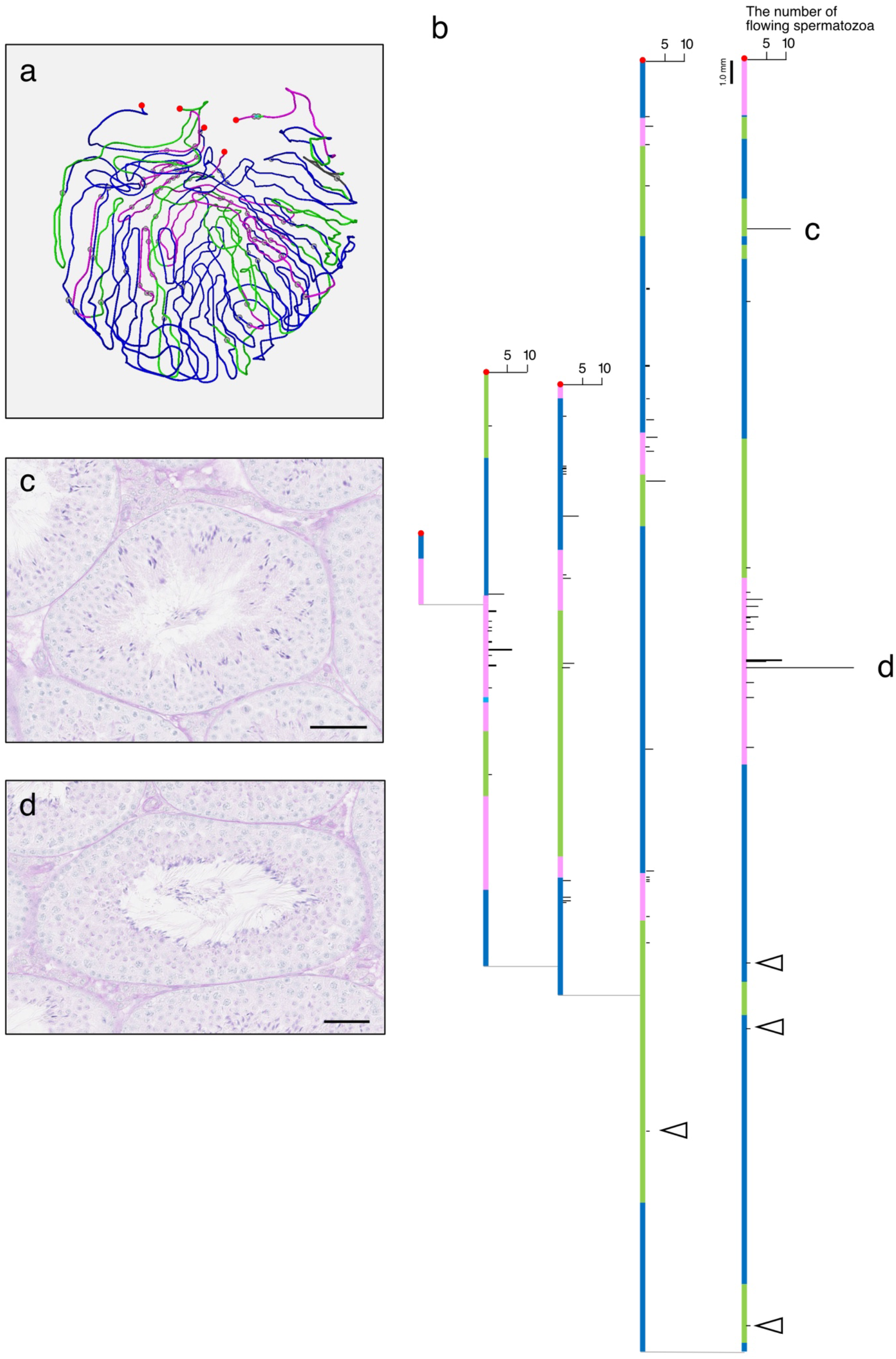
Representative localization of flowing spermatozoa in a seminiferous tubule (#2-ST5). (a) A reconstructed seminiferous tubule in a caudal view. Red points indicate the connection to the rete testis. Each node indicates the position of flowing spermatozoa. Each color represents the stage of the seminiferous epithelium (green: stage I-VI, magenta: stage VII/VIII, blue: stage IX-XII). (b) Histogram of the number of the flowing spermatozoa in a reconstructed seminiferous tubule. Each color bar represents the length of the seminiferous tubule at each stage (green: stage I-VI, magenta: stage VII/VIII, blue: stage IX-XII). Red points represent the rete testis, and each narrow horizontal line connecting each bar represents the branching point. Arrowheads indicate the flowing spermatozoa in the opposite direction to the rete testis. (c,d) Representative pictures of clustered spermatozoa in the seminiferous tubule. Bars = 50 μm.

In every testis and reconstructed ST, more than half of the flowing spermatozoa were found at stage VII/VIII (Tables 1,2). To clarify their localization in an ST, the distance of the flowing spermatozoa from the rete testis was recorded. Although there were more flowing spermatozoa at stage VII/VIII, they were scattered in the ST (Fig. 2b). On the other hand, clustered spermatozoa were also observed in one or two regions from each of the six STs (Fig. 2c,d). At first, we speculated that the flowing spermatozoa released from some seminiferous epithelium at stage VIII are accumulated toward the rete testis in an ST, and therefore that the flowing spermatozoa are found more frequently near the rete testis. However, the localization of the flowing spermatozoa was not biased through each ST, and therefore the increased accumulation of the flowing spermatozoa toward the rete testis was not observed (Fig. 2b). Furthermore, the flowing spermatozoa were also found in the opposite direction to the rete testis (Fig. 2b).

**Table 2.** The number of counted spermatozoa in each seminiferous tubule

|  |  | Spem count |  | Length (mm) |  | Sperm count<br>(/mm) |  |
| --- | --- | --- | --- | --- | --- | --- | --- |
|  | ST2 | 138 | (59) | 89.8 | (11.8) | 1.5 | (5.0) |
| #1 | ST6 | 113 | (86) | 106.4 | (22.9) | 1.1 | (3.8) |
|  | ST9 | 209 | (103) | 118 | (23.1) | 1.8 | (4.5) |
|  | ST2 | 35 | (17) | 91.9 | (6.2) | 0.4 | (2.8) |
| #2 | ST5 | 207 | (137) | 169.7 | (28.4) | 1.2 | (4.8) |
|  | ST10 | 187 | (105) | 109.7 | (17.3) | 1.7 | (6.1) |
|  | ST2 | 442 | (291) | 223.8 | (7.7) | 2.0 | (37.8) |
| #3 | ST5 | 342 | (238) | 108.8 | (10.8) | 3.1 | (10.8) |
|  | ST10 | 143 | (92) | 24.6 | (17.7) | 5.8 | (17.7) |

Values in stage VII/VIII are represented in the parenthesis

### *In vivo* imaging

Based on the above results, we hypothesized that the luminal flow could be changeable in an ST. Therefore, we tried to observe the luminal movement by *in vivo* imaging using a two-photon excitation microscope system. A fluorescence reporter mouse line, which is called Pax2-LynVenus and expresses a yellow fluorescent protein, Venus, on the cell membrane of the sperm flagellum, was created (Fig. 3). An anesthetized reporter mouse was placed on the warmed stage of the microscope after its testis was pulled out from the scrotum (Fig. 4a). The cell membrane of germ cells after meiosis and flagella of spermatozoa attaching to the seminiferous epithelium in the STs showed positive for Venus (Fig. 4b-j). In the time-lapse imaging, the change of the flagella direction could be observed, and it was found that the flagella direction was repeatedly reversed in the ST. The direction was changed within a few seconds to a few tens of seconds, although the period between the reversal was not the same among observations (Fig. 4 and Video 1).

**Figure 3.**
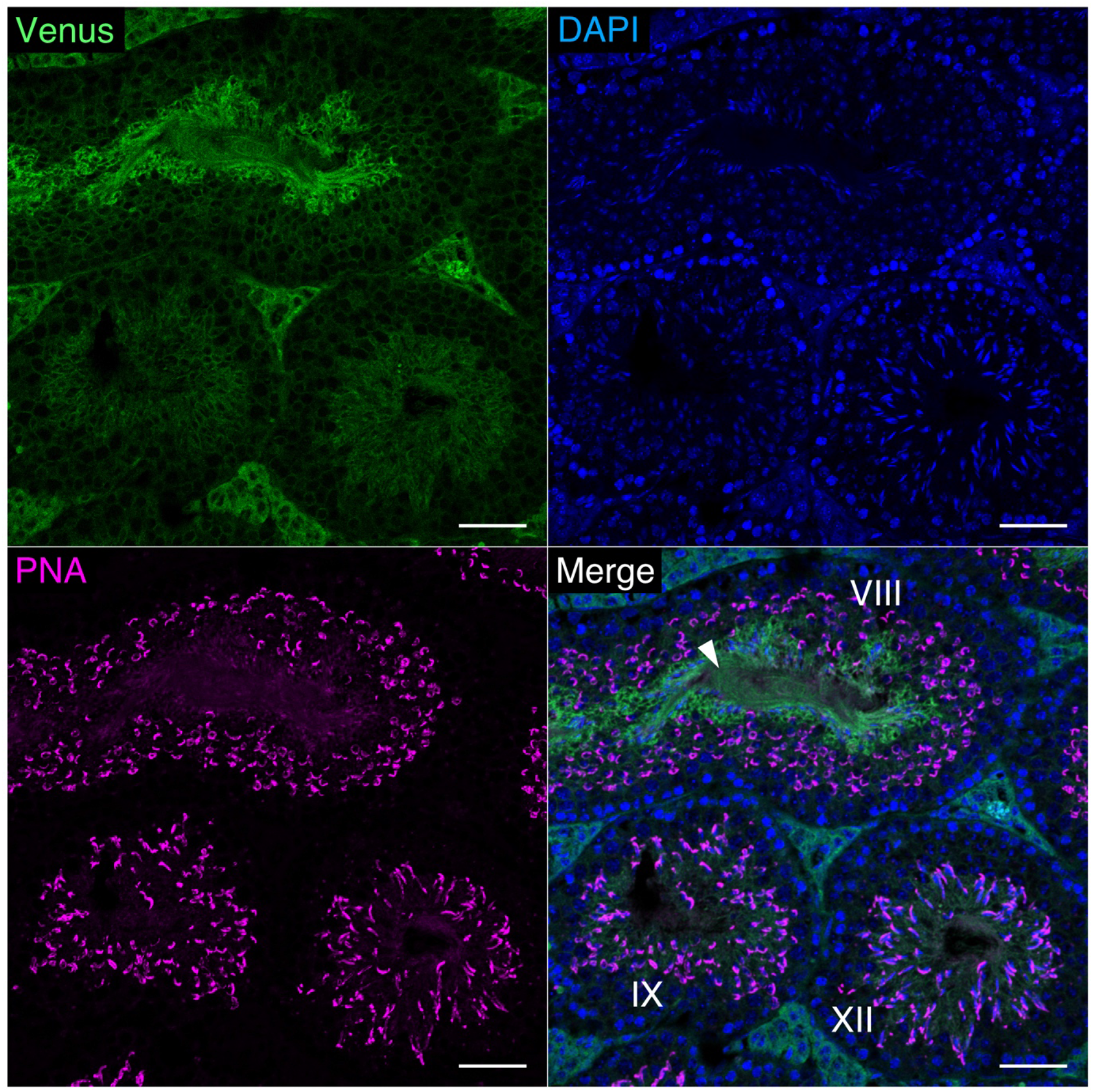
Representative pictures of the seminiferous tubules in the Pax2-LynVenus reporter mouse. Venus fluorescence is found on the cell membrane of germ cells, including flagella of the spermatozoa (an arrowhead), in the seminiferous tubules. Bars = 50 μm.

**Figure 4.**
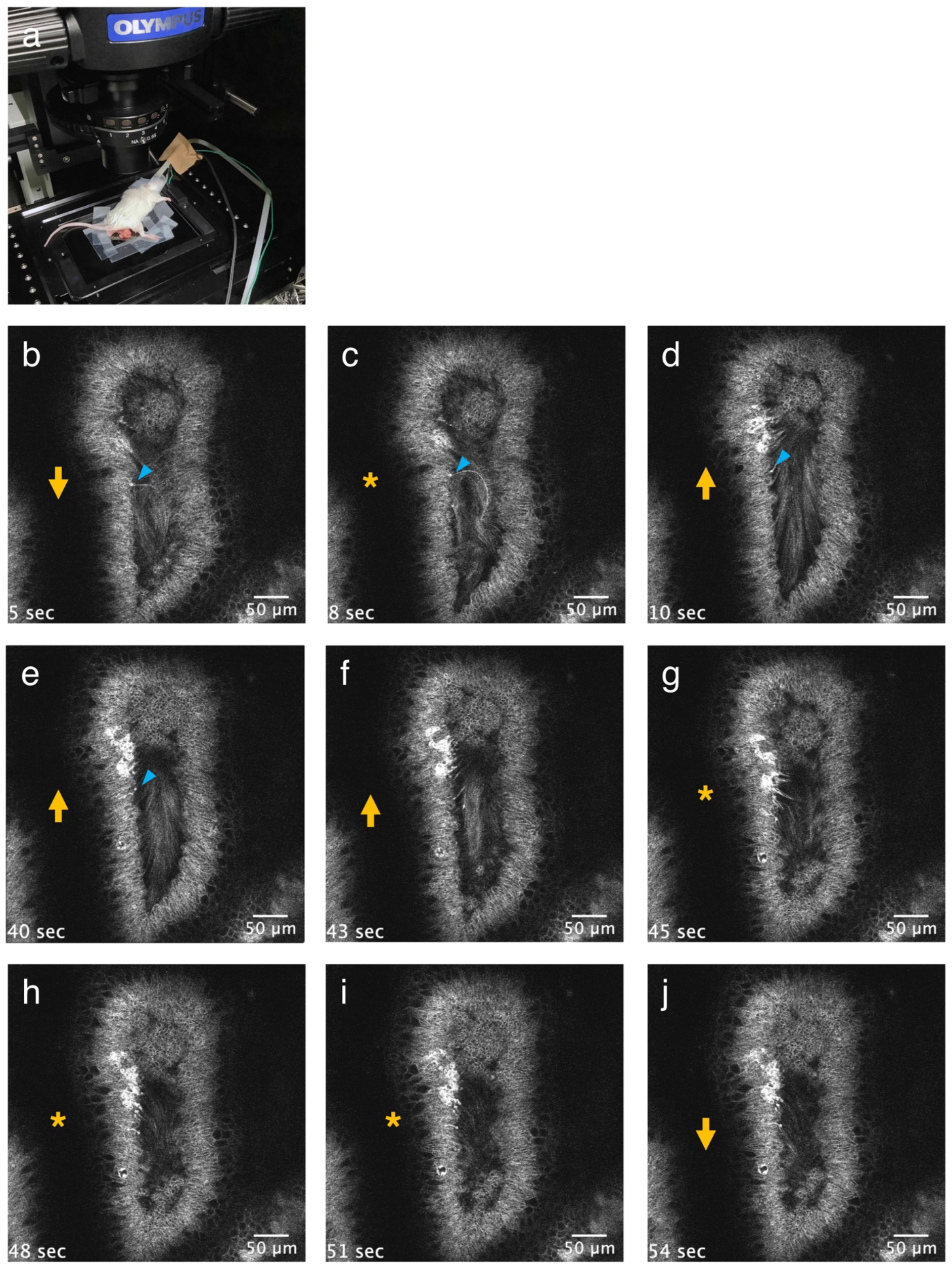
Representative time-lapse imaging of a seminiferous tubule at 2.7-second intervals. (a) A mouse is placed on the warmed stage with the testis withdrawn from the scrotum in a prone position during the observation. (b-j) The direction of flagella on the seminiferous epithelium is repeatedly reversed during the observation. Orange arrows and asterisks indicate the direction (b,d-f,j) and the transitional period (c,g-i), respectively. Bars = 50 μm.

We next focused on the curved region of the STs because the number of the spermatozoa attached to the seminiferous epithelium were different between the inner and outer curves (Fig. 1f). The curved ST at stage VII/VIII with the obvious sperm flagella was observed in the *in vivo* imaging. In the *in vivo* imaging, few flagella of the spermatozoa were also observed on the top of the seminiferous epithelium at the inner curve, resulting in the vacant space in the lumen at the inner side (Fig. 5 and Video 2). The flagella’s direction was also reversed in the curved region, and germ cells in the seminiferous epithelium also moved together with the flagella movements during the observation. Strikingly, the movements were more active in the inner curve than the outer one (Fig. 5 and Video 2).

**Figure 5.**
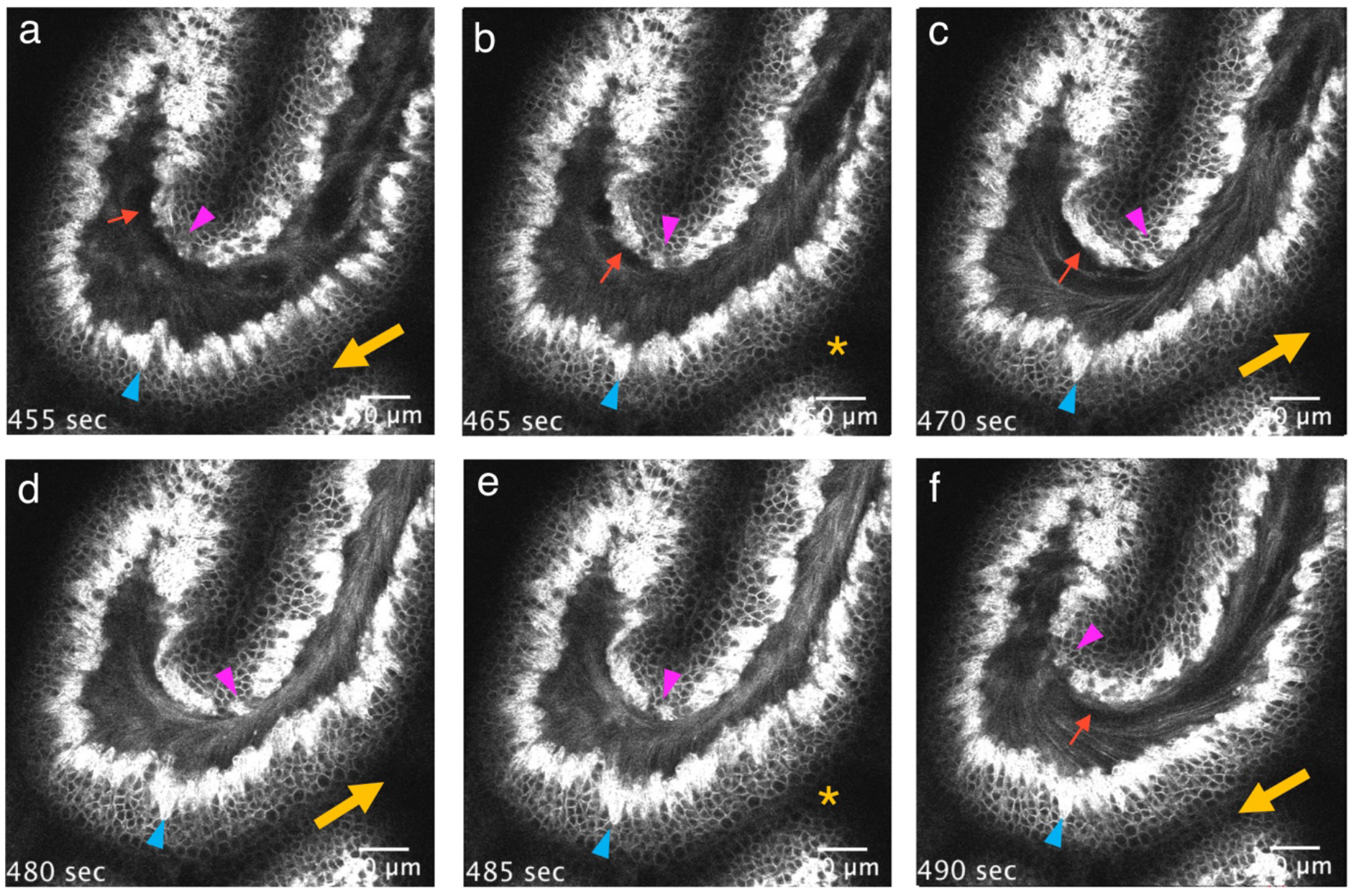
Representative time-lapse imaging of the curved seminiferous tubule at 4.9-second intervals. The direction of flagella is repeatedly reversed during the observation. Orange arrows indicate the direction (a,c,d,f), and the transitional period (asterisks) is shown in panels b and e. There are areas with few flagella at the inner side (red arrows). The position of the germ cells in the seminiferous epithelium is moved according to the luminal flow, and its movement is more extensive in the inner curve (magenta arrowheads) than the outer curve (blue arrowheads). Bars = 50 μm.

## Discussion

The present study investigated the luminal flow in the ST by three-dimensional analysis and *in vivo* imaging method. Investigating the flowing spermatozoa’s localization in a serial ST revealed that some flowing spermatozoa were present in the opposite direction to the rete testis. Besides, flagella movement was repeatedly reversed when observed by timelapse *in vivo* imaging. These results suggest that the luminal flow can be changed in the ST.

The spermatozoa are thought to be transported by the peristaltic motion of the ST to approach the epididymis. Disturbance of the PMC development induces dilation of the rete testis after puberty (Uchida *et al*., 2020). The flow of spermatozoa in the STs was recently observed by brightfield timelapse *in vivo* imaging (Fleck *et al*., 2021). They found that the cluster of the spermatozoa was transported in the ST in a unidirectional movement. On the other hand, *in vivo* imaging of the ST in the present study obtained bidirectional flagella movement, which was repeatedly reversed in the short term. This discrepancy is possibly due to the observation method and viewing scale. Imaging of the present study was a narrow region using two-photon excitation microscopy in contrast to the previous research that observed a vast area in brightfield. Therefore, they could observe macroscopic transport of the spermatozoa to the rete testis, and our results showed local luminal movements. The reversed flow may be induced by contraction and relaxation switching by the peritubular myoid cells.

Flowing spermatozoa were observed even in the opposite direction to the rete testis by three-dimensional analysis. This observation suggests that the liquid flow in the ST is not in a constant direction. Of course, all the spermatozoa will finally reach the rete testis. Therefore, the spermatozoa found in such a distal region would also be transported to the rete testis. A possible mechanism to pull spermatozoa to the rete testis is the water pressure difference between the STs and rete testis. Because more than 90% of the luminal fluid is absorbed into the epithelial cells in the efferent tubules (Clulow *et al*., 1994), luminal fluid with high pressure in the STs is probably pulled to the rete testis and efferent duct with low pressure of the luminal fluid. Further investigation of what induces the gradient of the luminal fluid pressure through the ST is necessary.

Fleck et al. (2021) observed agglomerated spermatozoa as transporting ones, and such clusters were also observed at one or two regions in an ST in this study. However, flowing spermatozoa were mainly scattered and observed more at stage Vll/Vlll than any other stages in the present histological analysis. We also observed bidirectional movements of the sperm flagella by *in vivo* imaging. From these observations, these flowing spermatozoa at stage VII/VIII are probably just after release from the Sertoli cell. The released spermatozoa will be aggregated by the bidirectional luminal flow and transported as a cluster.

In the present study, *in vivo* imaging showed that the seminiferous epithelium was repeatedly shaken together with the flagella movements of the spermatozoa on the epithelium. Furthermore, the seminiferous epithelium at the inner curve was vigorously shaken compared to that of the outer side when observed by *in vivo* imaging. On such an inner curve, few spermatozoa were present at the top of the epithelium. These findings suggest that shaking the Sertoli cells probably contributes to spermatozoa release. Although the shaking force may be the strongest in the inner curve of the STs, the contractile activities of the PMCs probably help release the spermatozoa even in the straight region.

Alternatively, the Sertoli cell’s ability to adhere to the spermatozoa may differ between the outer and inner curves. In our observation, spermatids at step 8 were found on the seminiferous epithelium, even at the inner curve. Although the spermatids after step 8 are assumed to be attached tightly to the Sertoli cells by the testis-specific adherens junctions, called an ectoplasmic specialization (Yan Cheng and Mruk, 2015), the spermatids are probably released between step 9–16 at the inner curve. From the results obtained in the present study, the shear force to the Sertoli cells must be different between the outer and inner curves. The cells subjected to the shear force react to resist the force by actin polymerization to strengthen their cytoplasm (Lee *et al*., 2006). The actin filament is tightly associated with the ectoplasmic specialization (Lie *et al*., 2010). Furthermore, the tubulobulbar complex emerges between the head of the spermatid and Sertoli cell at step 18–19 in the rat (Lie *et al*., 2010). The complex is a protrusion of the spermatid cytoplasm into the Sertoli cell and is tightly surrounded by the actin network in the Sertoli cell, although the precise function of the tubulobulbar complex is not known. Therefore, actin remodeling is necessary for both the ectoplasmic specialization and tubulobulbar complex during spermiogenesis. The different strength of the actin filament may cause malfunction of the adherent structures between the Sertoli cells and spermatozoa, resulting in the earlier release of the spermatozoa.

The present study is the first report that strongly suggests the contribution of PMCs to spermatozoa release. Of course, because these early spermatozoa released from the inner curve may not fully complete the spermiogenesis, they may possess relatively low fertility. This point leads to a hypothesis that the more active movements of the PMCs or external pressure may impair fertility due to the earlier release of the spermatozoa even after the emergence of the ectoplasmic specialization. The present study provides a new possibility that influence on the luminal flow can affect fertility in the male.

In conclusion, we could observe the transport of the spermatozoa throughout a single ST by 3D analysis and the movements in the STs by *in vivo* imaging using the Pax2-Lyn-Venus reporter mouse. The luminal flow is repeatedly reversed, and the physical force by the luminal flow probably leads to the release of the spermatozoa from the seminiferous epithelium. These results contribute to comprehending the release and transport of the spermatozoa in the STs.

## Author contribution statement

YK, TO, and HN performed the histological and three-dimensional analysis. TH created the Pax2-LynVenus reporter mouse and conducted *in vivo* imaging. All authors wrote and reviewed the manuscript.

## Acknowledgments

We are deeply grateful to the staff for animal care, especially to Katsuko Sudo, for her in vitro fertilization technique when introducing the Pax2-LynVenus mouse into Tokyo Medical University. We appreciate Professor Tatsunori Seki of the Department of Histology and Neuroanatomy, Tokyo Medical University, for the use of LSM700. We also appreciate Mr. Shuichi Yamazaki for his technical assistance in making serial paraffin sections. We thank Ms. Xi Wu, Ms. Miyuki Kuramasu, and Ms. Yuki Ogawa for their assistance in the routine procedures of histology and secretary works.

This work was supported by JSPS KAKENHI Grant Number JP16H06280, Grant-in-Aid for Scientific Research on Innovative Areas — Platforms for Advanced Technologies and Research Resources “Advanced Bioimaging Support.”

---

Video 1. Representative time-lapse imaging of a seminiferous tubule at 2.7-second intervals. The direction of flagella on the seminiferous epithelium is repeatedly reversed during the observation.

Video 2. Representative time-lapse imaging of the curved seminiferous tubule at 4.9-second intervals. The direction of flagella on the seminiferous epithelium is repeatedly reversed during the observation. The epithelium at the inner curve of the seminiferous tubule moved more actively and attached fewer spermatozoa compared to that at the outer curve.

